## supplemental figures for "Maternal obesity may disrupt offspring metabolism by inducing oocyte genome hyper-methylation via increased DNMTs"

### Supplementary Figures

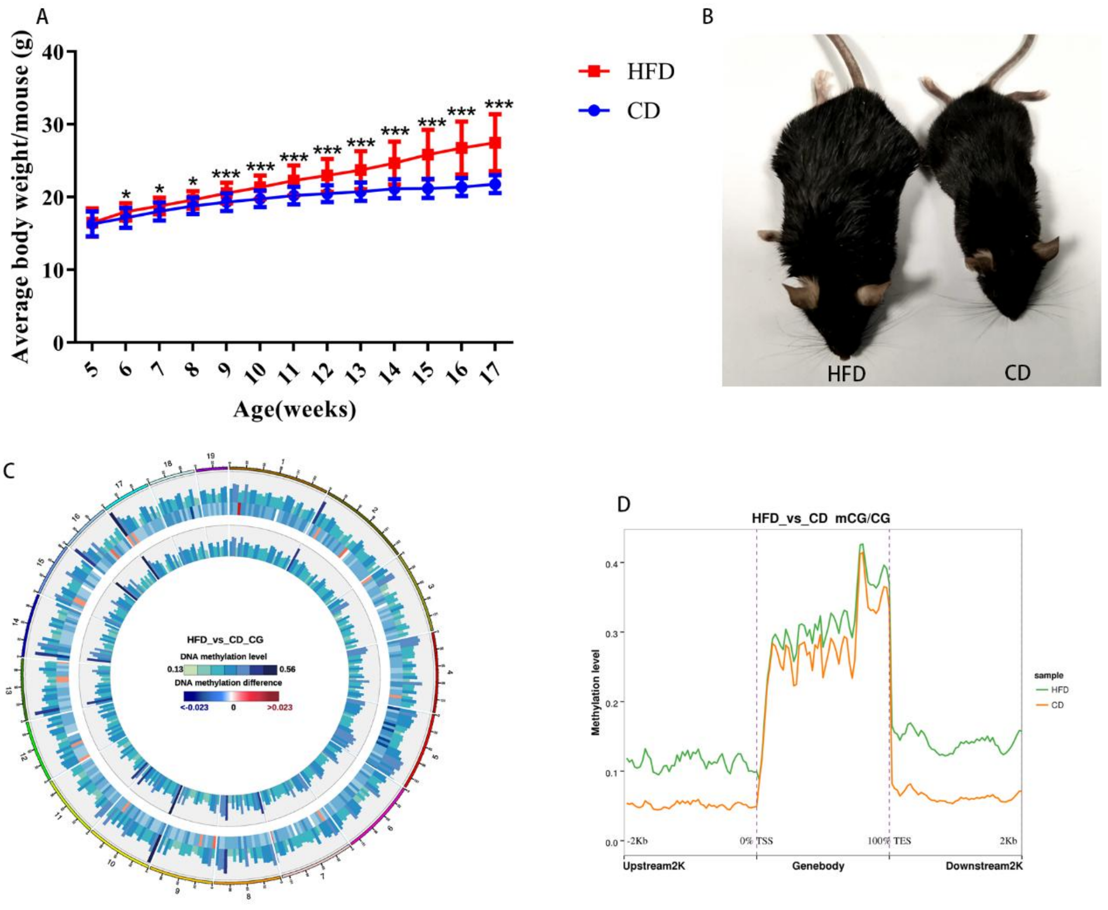

**Figure S1 Obese mouse model and DNA methylation in oocytes**
(A) Body weights of mice were examined every week.

(B) Mice fed with high-fat diet were obese compared with the control.

(C) Methylation level of CG distributed on chromosomes. From outside to inside circles: HFD methylation level, methylation difference between groups, control methylation level; color grading diagram, DNA methylation level; heat map, methylation difference level.

(D) Methylation distribution at different regions of genes.

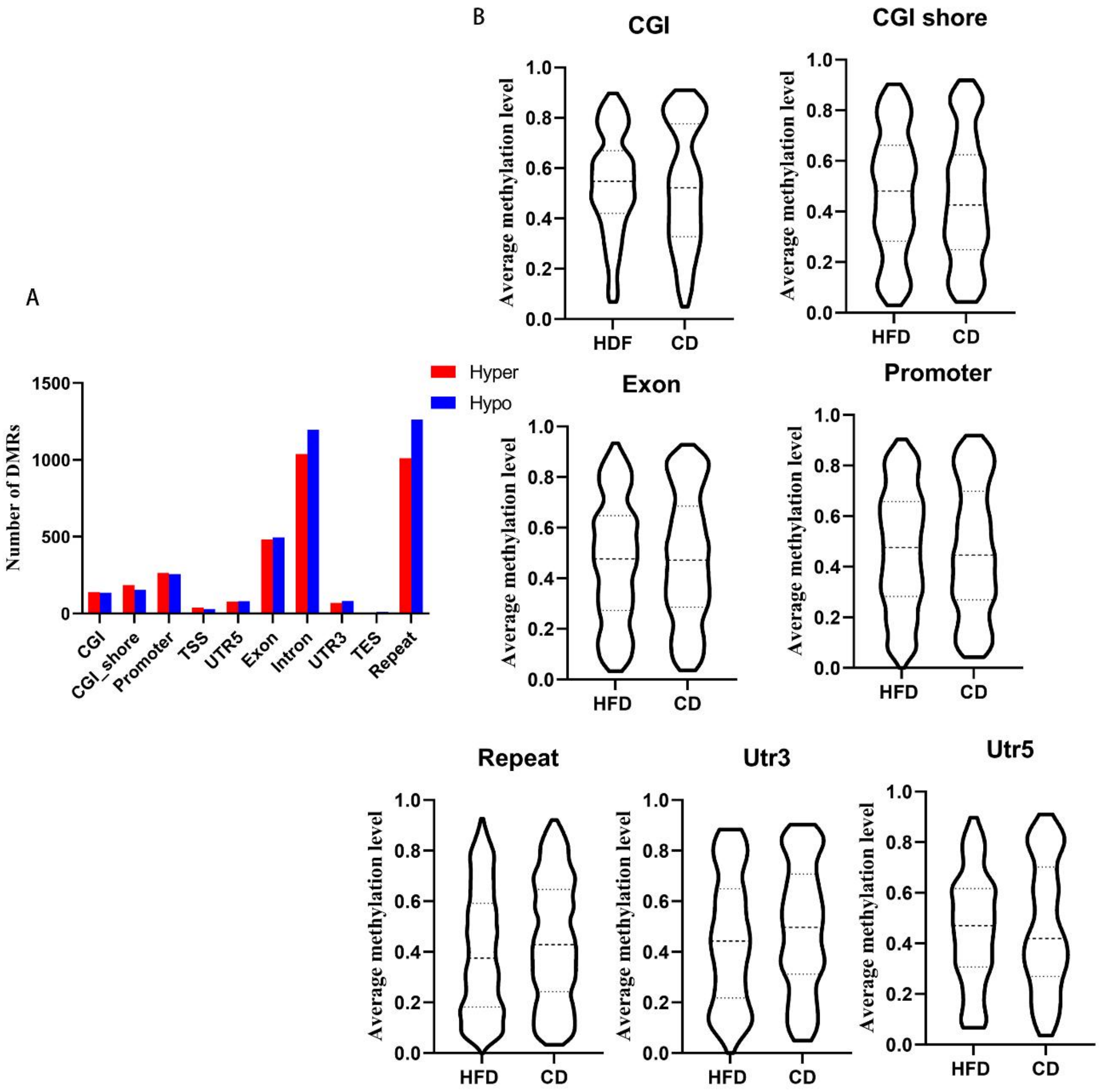


**Figure S2 DMRs methylation at different regions**

(A) DMRs numbers distribution at different elements.

(B) Average methylation levels of DMRs at elements.

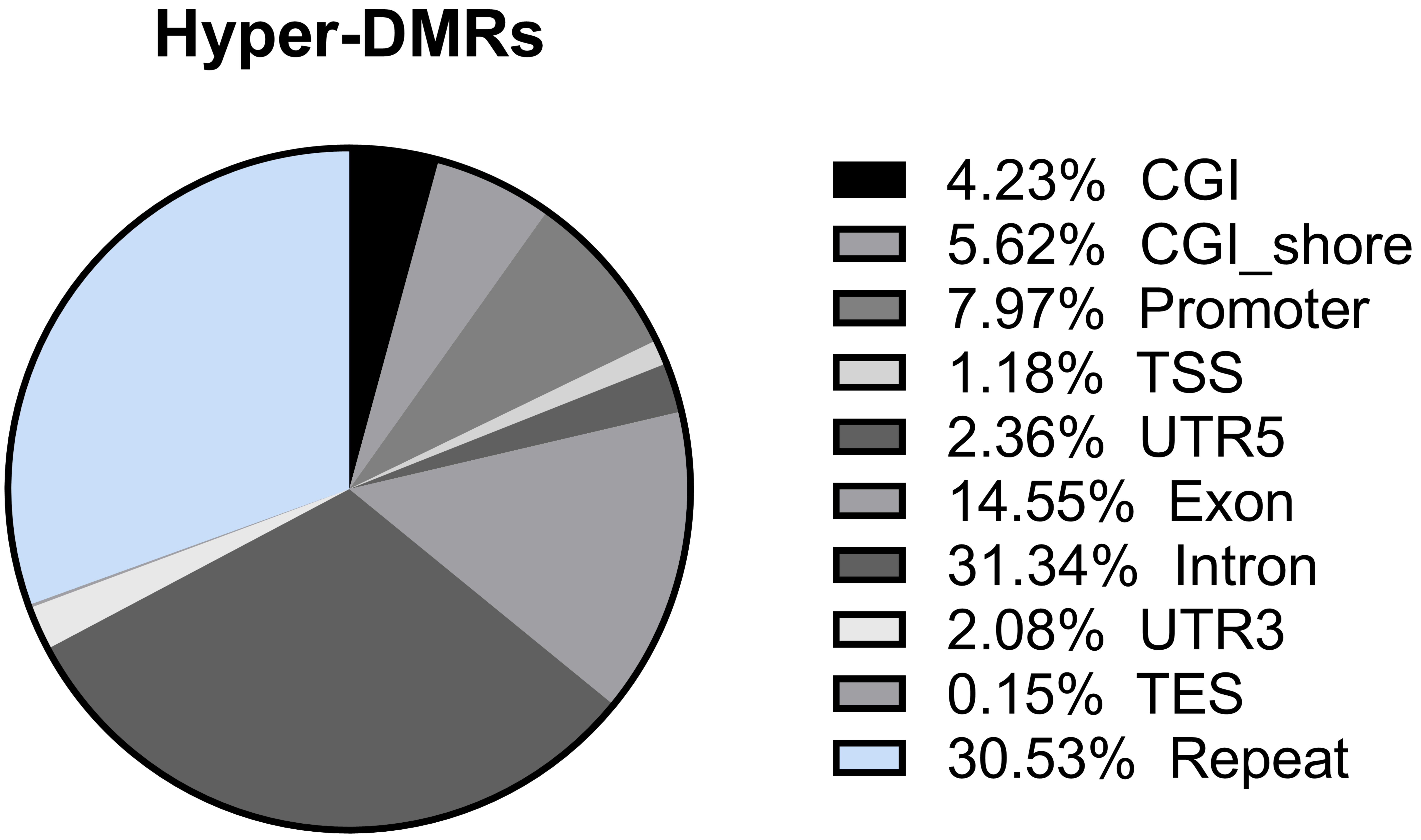
Hyper-DMRs
4.23% CGI
5.62% CGI_shore
7.97% Promoter
1.18% TSS
2.36% UTR5
14.55% Exon
31.34% Intron
2.08% UTR3
0.15% TES
30.53% Repeat

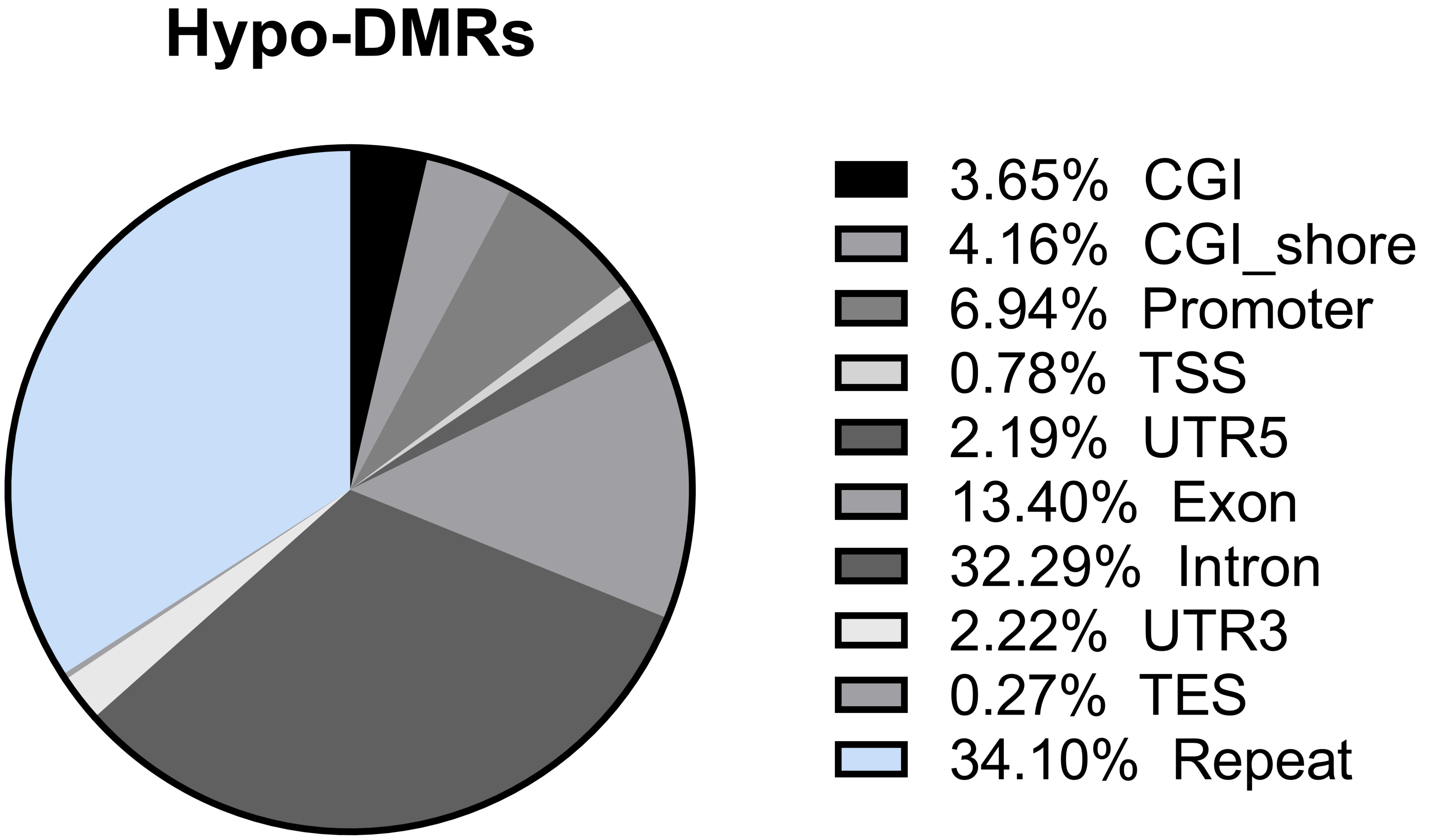
Hypo-DMRs
3.65% CGI
4.16% CGI_shore
6.94% Promoter
0.78% TSS
2.19% UTR5
13.40% Exon
32.29% Intron
2.22% UTR3
0.27% TES
34.10% Repeat

**Figure S3 distribution of the hypo- and hyper-DMRs in genomic elements.**

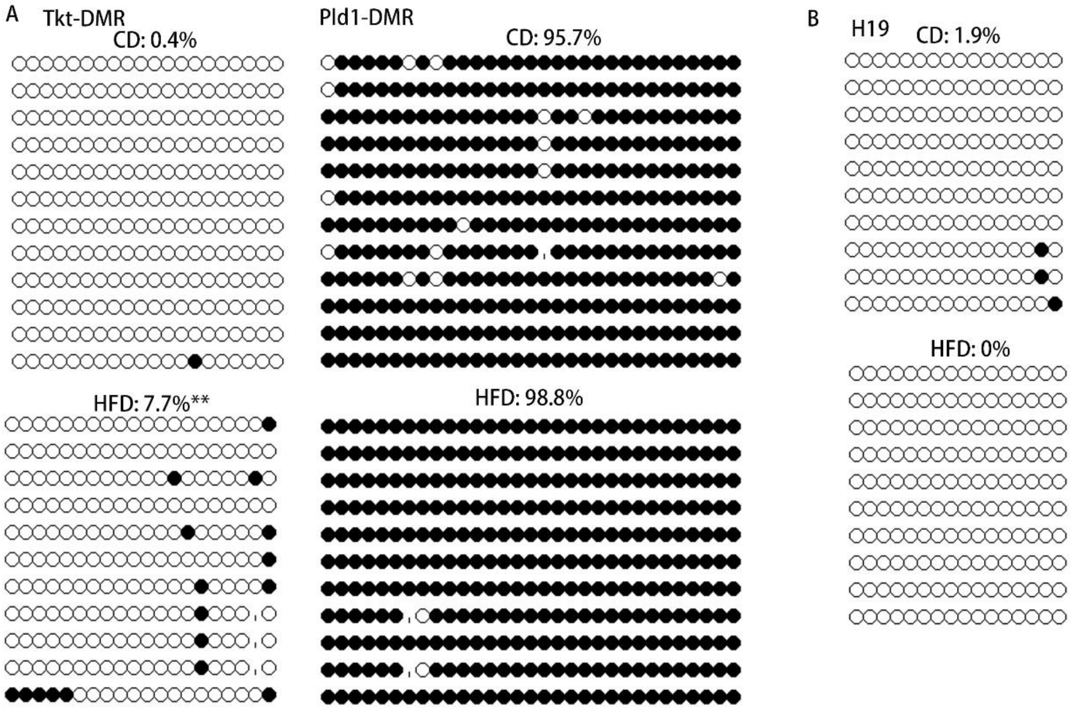


**Figure S4 Methylation of H19 in oocytes**

Methylation level of H19 in oocytes was examined using bisulfite sequencing. At least 10 available clones were used to calculate the methylation level, and total of 80-100 oocytes were analyzed for each group.

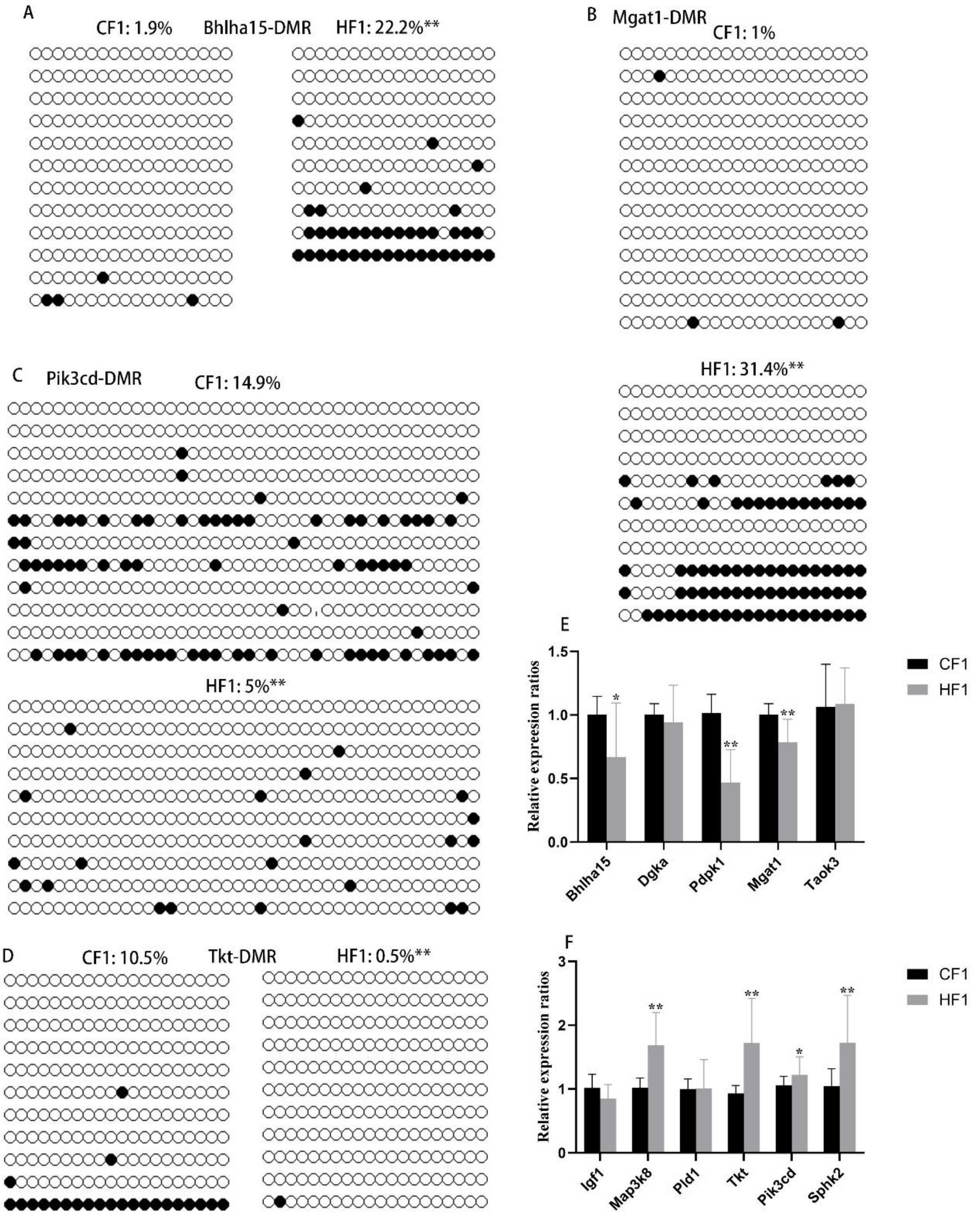
A
CF1: 1.9%
Bhlha15-DMR
HF1: 22.2%**
B
Mgat1-DMR
CF1: 1%
HF1: 31.4%**
C
Pik3cd-DMR
CF1: 14.9%
HF1: 5%**
D
CF1: 10.5%
Tkt-DMR
HF1: 0.5%**
E
Relative expression ratios
1.5
1.0
0.5
0.0
Bhlha15
Dgka
Pdpk1
Mgat1
Taok3
CF1
HF1
F
Relative expression ratios
3
2
1
0
Igf1
Map3k8
Pld1
Tkt
Pik3cd
Sphk2
CF1
HF1

**Figure S5 Methylation status of DMRs in F1 livers.**

(A) DMR methylation status at the promoter region of *Bhlha15*. CF1, female control mated with normal male; HF1, female obese mice mated with normal male. White circle, unmethylated CG; black circle, methylated CG.

(B) DMR methylation status at the promoter region of *Mgat1*. White circle, unmethylated CG; black circle, methylated CG.

(C) DMR methylation status at the promoter region of *Pik3cd*. White circle, unmethylated CG; black circle, methylated CG.

(D) DMR methylation status at the promoter region of *Tkt*. White circle, unmethylated CG; black circle, methylated CG.

(E and F) Relative expression of genes with hyper- or hypo-DMRs at promoter regions. * $p<0.05$; ** $p<0.01$.

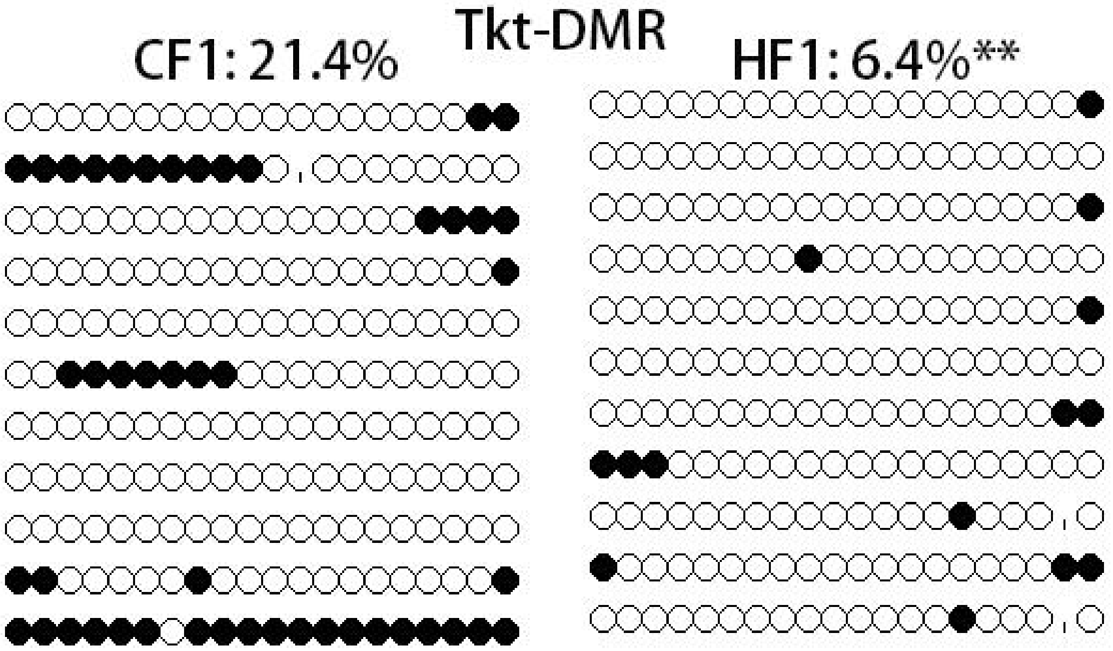


**Figure S6 Methylation level of Tkt-DMR in F1 oocytes.**

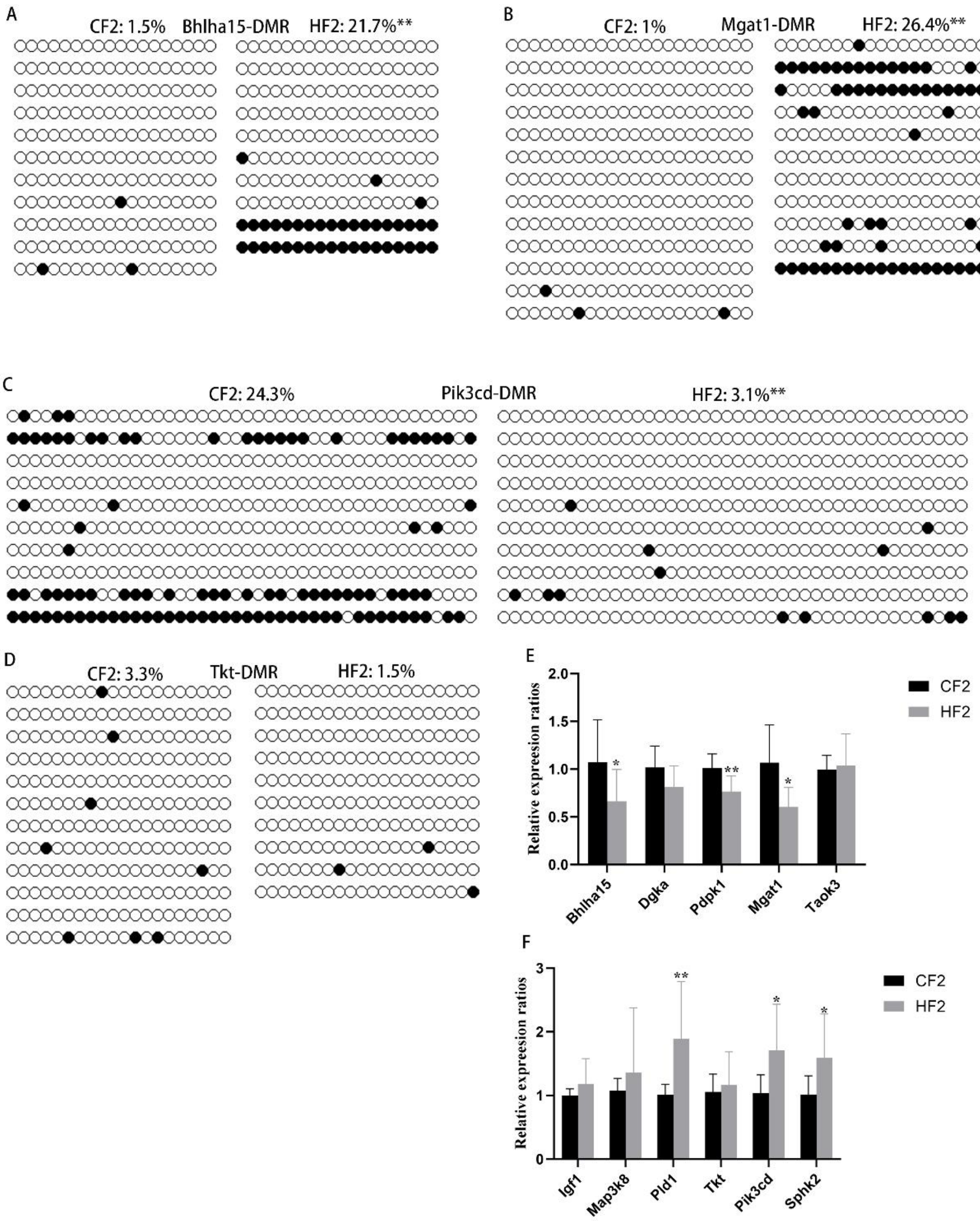
A
CF2: 1.5%
Bhlha15-DMR
HF2: 21.7%**
B
CF2: 1%
Mgat1-DMR
HF2: 26.4%**
C
CF2: 24.3%
Pik3cd-DMR
HF2: 3.1%**
D
CF2: 3.3%
Tkt-DMR
HF2: 1.5%
E
Relative expression ratios
2.0
1.5
1.0
0.5
0.0
*
**
*
Bhlha15
Dgka
Pdpk1
Mgat1
Taok3
CF2
HF2
F
Relative expression ratios
3
2
1
0
**
*
*
Igf1
Map3k8
Pld1
Tkt
Pik3cd
Sphk2
CF2
HF2

**Figure S7 Methylation of DMRs in F2 livers.**

(A-D) Methylation levels of *Bhlha15*-DMR, *Mgat1*-DMR, *Pik3cd*-DMR, and *Tkt*-DMR were respectively examined using bisulfite sequencing, and at least 10 available clones were used for each DMR. White circle, unmethylated CG; black circle, methylated CG. ** $p<0.01$.

(E) Expression of genes with hyper-DMRs at promoter regions in F2 livers. * $p<0.05$; ** $p<0.01$.

(F) Expression of genes with hypo-DMRs at promoter regions in F2 livers. * $p<0.05$; ** $p<0.01$.

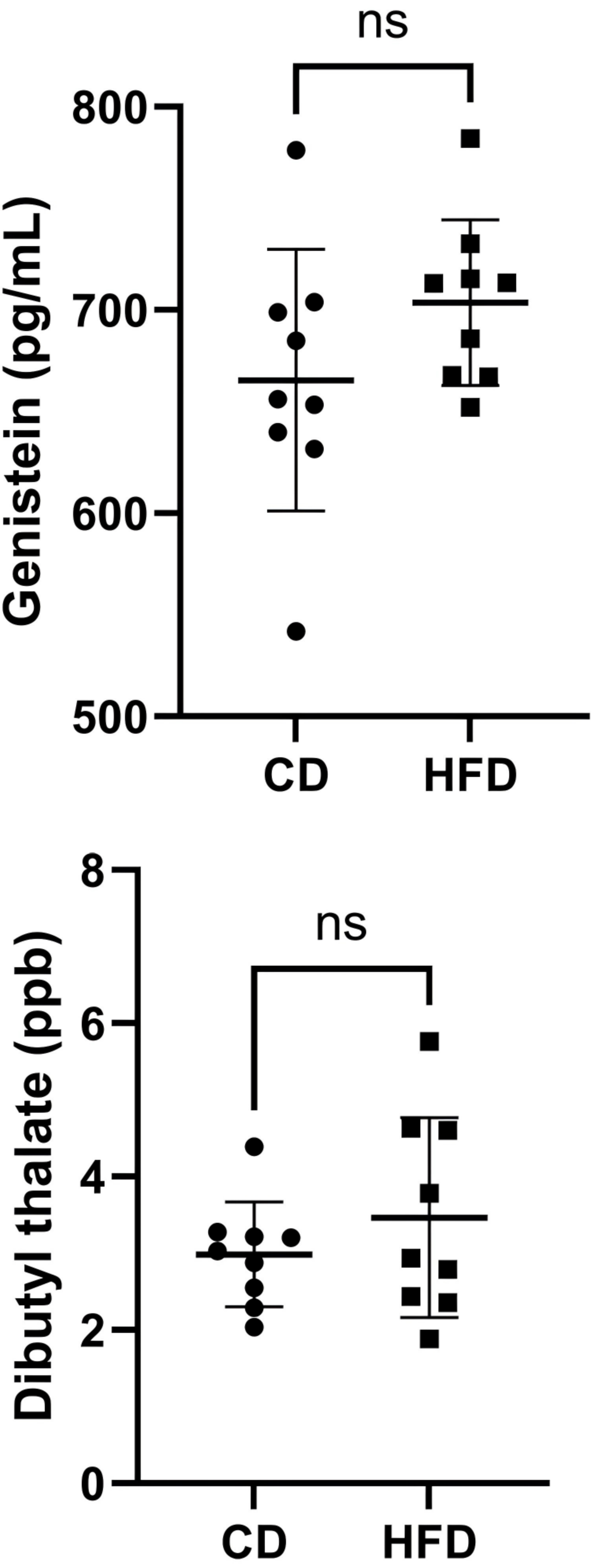
ns
800
700
600
500
Genistein (pg/mL)
CD
HFD
ns
8
6
4
2
0
Dibutyl thalate (ppb)
CD
HFD

**Figure S8 Concentrations of genistein and dibutylthalate in the seru m of HFD and CD.**

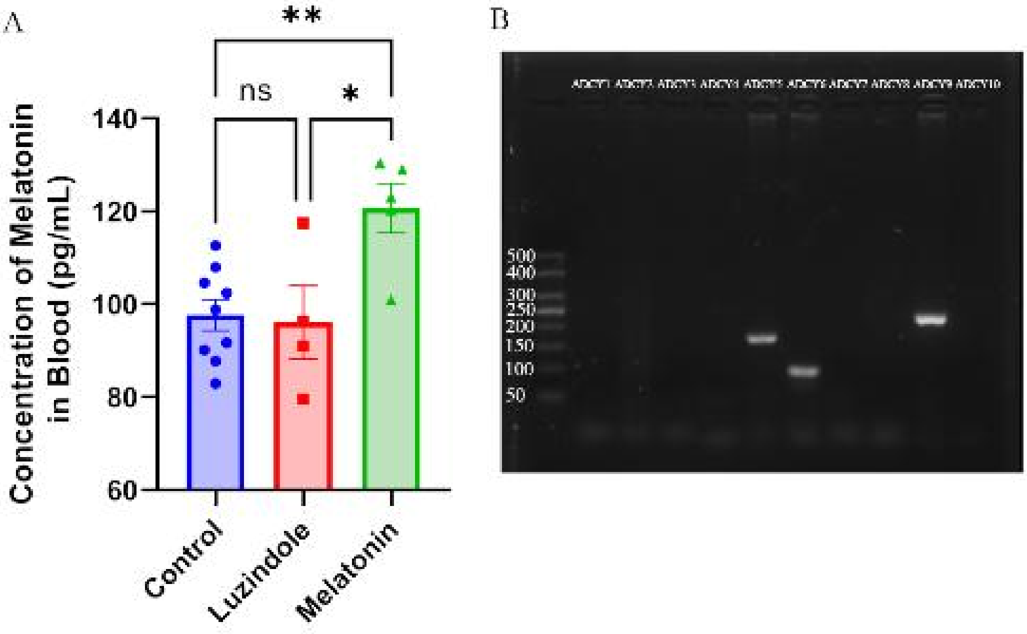


**Figure S9 Melatonin level in blood and ADCYs expression in oocytes**.

(A) Melatonin levels in blood of control mice, luzindole-treated mice, and exogenous melatonin-treated mice were examined using ELISA. * p<0.05; ** p<0.01.

(B) The expression of ADCYs in oocytes was examined using RT-PCR.

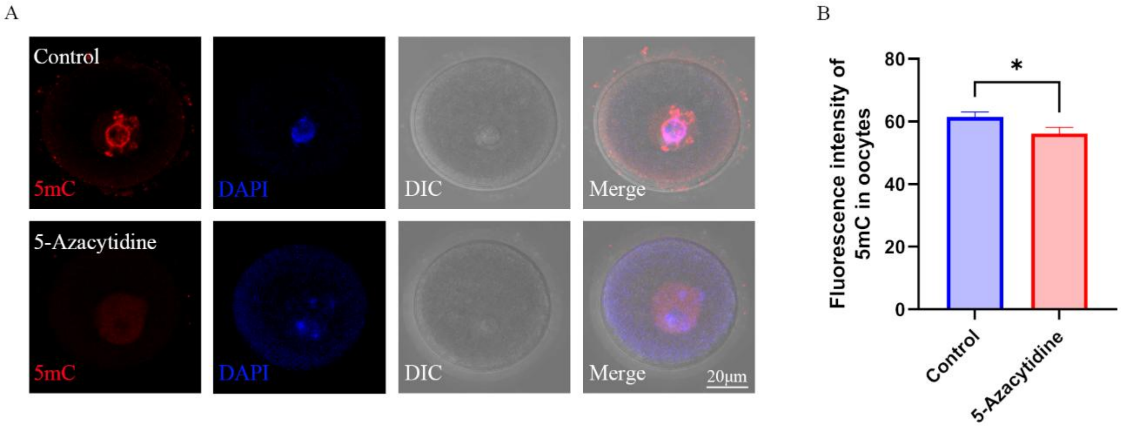


**Figure S10 DNMTs regulates DNA methylation in oocytes.**

(A) Female mice were treated with DNMTs inhibitor 5-azacytidine, and methylation in oocytes was examined using immunofluorescence.

(B) The relative fluorescence intensity was calculated using Image J. * $p<0.05$.
