## supplemental tables for "Maternal obesity may disrupt offspring metabolism by inducing oocyte genome hyper-methylation via increased DNMTs"

**Table S1 Some information of whole genome bisulfite sequencing.**

| Samples | Total reads | Mapped reads | Unique Mapping rate (%) | Mean Sequencing Depth | Conversion Rate (%) |
|---|---|---|---|---|---|
| CD1 | 102971225 | 33856938 | 32.88 | 2.46 | 99.449 |
| CD2 | 99413634 | 37359643 | 37.58 | 2.69 | 99.458 |
| HFD1 | 87762792 | 32867165 | 37.45 | 2.48 | 99.477 |
| HFD2 | 85697050 | 31613641 | 36.89 | 2.45 | 99.489 |
| HFD3 | 104081620 | 35991424 | 34.58 | 2.75 | 99.501 |

**Table S2. KEGG pathway analysis of metabolism-relative genes**

| KEGG class | Pathway | Out (147) | All (9198) | Pathway ID | Genes |
|---|---|---|---|---|---|
| Amino acid metabolism | Glycine, serine and threonine metabolism | 2 | 42 | ko00260 | ENSMUSG00000017713;ENSMUSG00000024039 |
| Amino acid metabolism | Tyrosine metabolism | 1 | 40 | ko00350 | ENSMUSG00000055301 |
| Amino acid metabolism | Cysteine and methionine metabolism | 1 | 54 | ko00270 | ENSMUSG00000024039 |
| Amino acid metabolism | Valine, leucine and isoleucine degradation | 1 | 57 | ko00280 | ENSMUSG00000062908 |
| Amino acid metabolism | Lysine degradation | 1 | 66 | ko00310 | ENSMUSG00000002028 |
| Carbohydrate metabolism | Inositol phosphate metabolism | 4 | 73 | ko00562 | ENSMUSG00000039936;ENSMUSG00000023805;ENSMUSG00000028894;ENSMUSG00000025477 |
| Carbohydrate metabolism | Amino sugar and nucleotide sugar metabolism | 2 | 51 | ko00520 | ENSMUSG00000031387;ENSMUSG00000028671 |
| Carbohydrate metabolism | Citrate cycle (TCA cycle) | 1 | 32 | ko00020 | ENSMUSG00000061838 |
| Carbohydrate metabolism | Pentose phosphate pathway | 1 | 33 | ko00030 | ENSMUSG00000021957 |
| Carbohydrate metabolism | Galactose metabolism | 1 | 34 | ko00052 | ENSMUSG00000028671 |
| Carbohydrate metabolism | Propanoate metabolism | 1 | 34 | ko00640 | ENSMUSG00000061838 |
| Carbohydrate metabolism | Glycolysis / Gluconeogenesis | 1 | 70 | ko00010 | ENSMUSG00000055301 |
| Energy metabolism | Nitrogen metabolism | 1 | 17 | ko00910 | ENSMUSG00000028463 |
| Energy metabolism | Oxidative phosphorylation | 1 | 125 | ko00190 | ENSMUSG00000022450 |

| | | | | | |
|---|---|---|---|---|---|
| Global and overview maps | Fatty acid metabolism | 3 | 61 | ko01212 | ENSMUSG00000062908;ENSMUSG00000007783;ENSMUSG00000025153 |
| Global and overview maps | Biosynthesis of amino acids | 3 | 81 | ko01230 | ENSMUSG00000021957;ENSMUSG00000017713;ENSMUSG00000024039 |
| Global and overview maps | Carbon metabolism | 2 | 124 | ko01200 | ENSMUSG00000021957;ENSMUSG00000061838 |
| Glycan biosynthesis and metabolism | Various types of N-glycan biosynthesis | 2 | 40 | ko00513 | ENSMUSG00000043998;ENSMUSG00000020346 |
| Glycan biosynthesis and metabolism | N-Glycan biosynthesis | 2 | 51 | ko00510 | ENSMUSG00000043998;ENSMUSG00000020346 |
| Glycan biosynthesis and metabolism | Other glycan degradation | 1 | 18 | ko00511 | ENSMUSG00000033857 |
| Glycan biosynthesis and metabolism | Glycosylphosphatidylinositol(GPI)-anchor biosynthesis | 1 | 25 | ko00563 | ENSMUSG00000014245 |
| Glycan biosynthesis and metabolism | Glycosphingolipid biosynthesis - lacto and neolacto series | 1 | 28 | ko00601 | ENSMUSG00000021360 |
| Lipid metabolism | Ether lipid metabolism | 4 | 48 | ko00565 | ENSMUSG00000030703;ENSMUSG00000023913;ENSMUSG00000027695;ENSMUSG00000049721 |
| Lipid metabolism | Sphingolipid metabolism | 3 | 47 | ko00600 | ENSMUSG00000021263;ENSMUSG00000057342;ENSMUSG00000049721 |
| Lipid metabolism | Fatty acid degradation | 3 | 52 | ko00071 | ENSMUSG00000062908;ENSMUSG00000055301;ENSMUSG00000007783 |

| | | | | | |
|---|---|---|---|---|---|
| Lipid metabolism | Fatty acid biosynthesis | 1 | 18 | ko00061 | ENSMUSG00000025153 |
| Lipid metabolism | Steroid biosynthesis | 1 | 22 | ko00100 | ENSMUSG00000024799 |
| Lipid metabolism | Glycerophospholipid metabolism | 2 | 101 | ko00564 | ENSMUSG00000025357;ENSMUSG00000027695 |
| Lipid metabolism | Glycerolipid metabolism | 1 | 65 | ko00561 | ENSMUSG00000025357 |
| Lipid metabolism | Steroid hormone biosynthesis | 1 | 103 | ko00140 | ENSMUSG00000024365 |
| Metabolism of cofactors and vitamins | Thiamine metabolism | 1 | 16 | ko00730 | ENSMUSG00000026807 |
| Metabolism of cofactors and vitamins | Porphyrin and chlorophyll metabolism | 1 | 43 | ko00860 | ENSMUSG00000028684 |
| Metabolism of cofactors and vitamins | Nicotinate and nicotinamide metabolism | 1 | 44 | ko00760 | ENSMUSG00000029063 |
| Metabolism of cofactors and vitamins | Retinol metabolism | 1 | 102 | ko00830 | ENSMUSG00000055301 |
| Metabolism of other amino acids | Glutathione metabolism | 1 | 68 | ko00480 | ENSMUSG00000022562 |
| Nucleotide metabolism | Purine metabolism | 4 | 139 | ko00230 | ENSMUSG00000021699;ENSMUSG00000026807;ENSMUSG00000021684;ENSMUSG00000042638 |
| Xenobiotics biodegradation and metabolism | Drug metabolism - cytochrome P450 | 1 | 74 | ko00982 | ENSMUSG00000055301 |
| Xenobiotics biodegradation and metabolism | Metabolism of xenobiotics by cytochrome | 1 | 83 | ko00980 | ENSMUSG00000055301 |

| | | | | | |
|---|---|---|---|---|---|
| | P450 | | | | |
| Xenobiotics biodegradation and metabolism | Drug metabolism - other enzymes | 1 | 97 | ko00983 | ENSMUSG00000009350 |

**Table S3. DMRs methylation status of metabolism-relative genes**

| Gene ID | Symbol | Hyper/Hypo |
|---|---|---|
| ENSMUSG00000014245 | Pigl | Hyper |
| ENSMUSG00000062908 | Acadm | Hyper |
| ENSMUSG00000055301 | Adh7 | Hyper |
| ENSMUSG00000020346 | Mgat1 | Hyper |
| ENSMUSG00000021263 | Degs2 | Hyper |
| ENSMUSG00000021360 | Gcnt2 | Hyper |
| ENSMUSG00000021699 | Pde4d | Hyper |
| ENSMUSG00000022450 | Ndufa6 | Hyper |
| ENSMUSG00000022562 | Oplah | Hyper |
| ENSMUSG00000023913 | Pla2g7 | Hyper |
| ENSMUSG00000024039 | Cbs | Hyper |
| ENSMUSG00000024365 | Cyp21a1 | Hyper |
| ENSMUSG00000024799 | Tm7sf2 | Hyper |
| ENSMUSG00000025357 | Dgka | Hyper |
| ENSMUSG00000028894 | Inpp5b | Hyper |
| ENSMUSG00000029063 | Nadk | Hyper |
| ENSMUSG00000028463 | Car9 | Hyper |
| ENSMUSG00000028671 | Gale | Hyper |
| ENSMUSG00000028684 | Urod | Hyper |
| ENSMUSG00000002028 | Kmt2a | Hypo |
| ENSMUSG00000017713 | Tha1 | Hypo |
| ENSMUSG00000049721 | Gal3st1 | Hypo |
| ENSMUSG00000057342 | Sphk2 | Hypo |
| ENSMUSG00000061838 | Suclg2 | Hypo |
| ENSMUSG00000042638 | Gucy2c | Hypo |
| ENSMUSG00000043998 | Mgat2 | Hypo |
| ENSMUSG00000039936 | Pik3cd | Hypo |
| ENSMUSG00000021684 | Pde8b | Hypo |
| ENSMUSG00000021957 | Tkt | Hypo |
| ENSMUSG00000026807 | Ak8 | Hypo |
| ENSMUSG00000023805 | Synj2 | Hypo |
| ENSMUSG00000025477 | Inpp5a | Hypo |
| ENSMUSG00000025153 | Fasn | Hypo |
| ENSMUSG00000027695 | Pld1 | Hypo |
| ENSMUSG00000031387 | Renbp | Hypo |

**Table S4. Binding sites of CREB1 on sequences of DNMTs**

| Name | Relative score | Sequence ID | Start | End | Strand | Predicted sequence |
|---|---|---|---|---|---|---|
| Chr12: 3976409:3977697 | | | | | | |
| MA0018.2.CREB1 | 0.8299 | DNMT3a | 431 | 438 | - | TGAGGCCT |
| MA0018.2.CREB1 | 0.8092 | DNMT3a | 42 | 49 | + | TGAGGTCC |
| MA0018.2.CREB1 | 0.8012 | DNMT3a | 1255 | 1262 | - | CGAGGCCA |
| 12:3875426:3875897 | | | | | | |
| MA0018.2.CREB1 | 0.8299 | DNMT3a | 372 | 379 | - | TGAGGCCT |
| Chr12: 3977963:3978629 | | | | | | |
| MA0018.2.CREB1 | 0.8901 | DNMT3a | 129 | 136 | + | TGAGGCCA |
| Chr9: 20864302:20865039 | | | | | | |
| MA0018.2.CREB1 | 0.9342 | DNMT1 | 408 | 415 | + | TGAGGTCA |
| MA0018.2.CREB1 | 0.8642 | DNMT1 | 408 | 415 | - | TGACCTCA |
| Chr10: 77897008:77897613 | | | | | | |
| MA0018.2.CREB1 | 0.8574 | DNMT3l | 336 | 343 | + | TGAGGTGA |
| MA0018.2.CREB1 | 0.8092 | DNMT3l | 507 | 514 | - | TGAGGGCA |
| MA0018.2.CREB1 | 0.8012 | DNMT3l | 405 | 412 | - | GGAGGCCA |

Note: Sequences are obtained from Cut & Tag assay. Chr: chromosome.

Table S5. Annotation of peaks at relative gene regions

| Peak | Chr | Start | End | Length | -Log10 (p value) | Annotation | Gene Chr | Gene Start | Gene End | Gene Length | SYMBOL |
|---|---|---|---|---|---|---|---|---|---|---|---|
| peak2_1593 | chr10 | 77897008 | 77897613 | 606 | 12.4365 | Intron (ENSMUST00000138785.8/54427, intron 13 of 13) | chr10 | 77891169 | 77899449 | 8281 | Dnmt3l |
| peak2_2951 | chr12 | 3976409 | 3977697 | 1289 | 10.9896 | Distal Intergenic | chr12 | 3857160 | 3963491 | 106332 | Dnmt3a |
| peak2_2952 | chr12 | 3977963 | 3978629 | 667 | 8.99516 | Distal Intergenic | chr12 | 3857160 | 3963491 | 106332 | Dnmt3a |
| peak2_415 | chr12 | 3875426 | 3875897 | 472 | 12.1207 | Intron (ENSMUST00000173700.2/13435, intron 1 of 1) | chr12 | 3857160 | 3963491 | 106332 | Dnmt3a |
| peak2_14239 | chr9 | 20864302 | 20865039 | 738 | 10.9407 | Promoter (<=1kb) | chr9 | 20818505 | 20864275 | 45771 | Dnmt1 |

Note: Chr, chromosome.

**Table S6 Percentage of ingredient of diets.**

| Class description | D12492 (%) | Normal Diet (%) |
|---|---|---|
| Protein | 26.23 | 24.02 |
| Carbohydrate | 25.56 | 29.75 |
| Fiber | 6.46 | ≤5 |
| Fat | 34.89 | 12.95 |
| Mineral | 6.46 | 4.42 |
| Vitamin | 0.39 | 0.36 |
| Dye | 0.0065 | 0 |
| H2O | | ≤10 |

Note: for D12492, the percentage is calculated using ingredient weight/773.85 (g) ×100%; for normal diet, the percentage is calculated using ingredient weight/1000 (g) ×100%.
